# The Caribbean-Hispanic Alzheimer’s Brain Transcriptome Reveals Ancestry-Specific Disease Mechanisms

**DOI:** 10.1101/2020.05.28.122234

**Authors:** Daniel Felsky, Sanjeev Sariya, Ismael Santa-Maria, Leon French, Julie A. Schneider, David A. Bennett, Richard Mayeux, Philip L. De Jager, Giuseppe Tosto

## Abstract

Ethnicity impacts Alzheimer’s disease risk, especially among Caribbean-Hispanics. We report the first RNA-sequencing analysis of brain tissue from 45 Alzheimer’s disease and control Caribbean-Hispanics. Data were compared with two independent samples of non-Hispanic Caucasians (total n=729). By identifying and characterizing those genes with ancestry- and region-specific expression patterns in Alzheimer’s disease, we reveal molecular insights that may help explain epidemiological disparities in this understudied aging population.

## Main text

Genetic and environmental factors conferring risk for late-onset Alzheimer’s disease (LOAD) are known to differ across ethnic groups,^1^ though genomic studies are still dominated by non-Hispanic White (NHW) cohorts,^2^ particularly for neurodegenerative diseases. Here we present the first brain gene expression study from Caribbean-Hispanics (CH) with and without LOAD. We aimed to identify genes and pathways with ancestry-specific (i.e. those identified in CH only) and ancestry-independent (i.e. those replicating across groups) differential gene expression by comparing transcriptome-wide association analyses (TWAS) and co-expression network based analyses performed in CH and in two independent NHW cohorts, processed with identical pipelines.

We performed RNA-sequencing and whole-genome genotyping on postmortem temporal cortex (TCX) tissue from 45 self-reported CH individuals ascertained from the New York Brain Bank (NYBB). Global admixture analysis^3^ revealed that seven individuals did not show all three ancestral components (European, African and Native-American) (**see Supplementary Information, Supplementary Figure 1**), resulting in 38 genetically-confirmed three-way admixed subjects (n_LOAD_=20, n_non-LOAD_=18); seven carried the *PSEN1* G206A mutation, a known mutation associated with familial AD in Caribbean Hispanic populations.^4^ Following data quality control (QC), robust regression modeling of LOAD status on gene transcript abundance (log_2_(CPM)) was adjusted for demographic and technical covariates (see **Supplementary Information**) and carried out in three stages: “stage 1”) including all (n=45), “stage 2”) excluding genetically non-CH (n=38), and “stage 3”) excluding both non-CH subjects and *PSEN1* carriers (n=31). Significance testing for differential expression was performed using empirical Bayes moderation.

**Figure 1.**
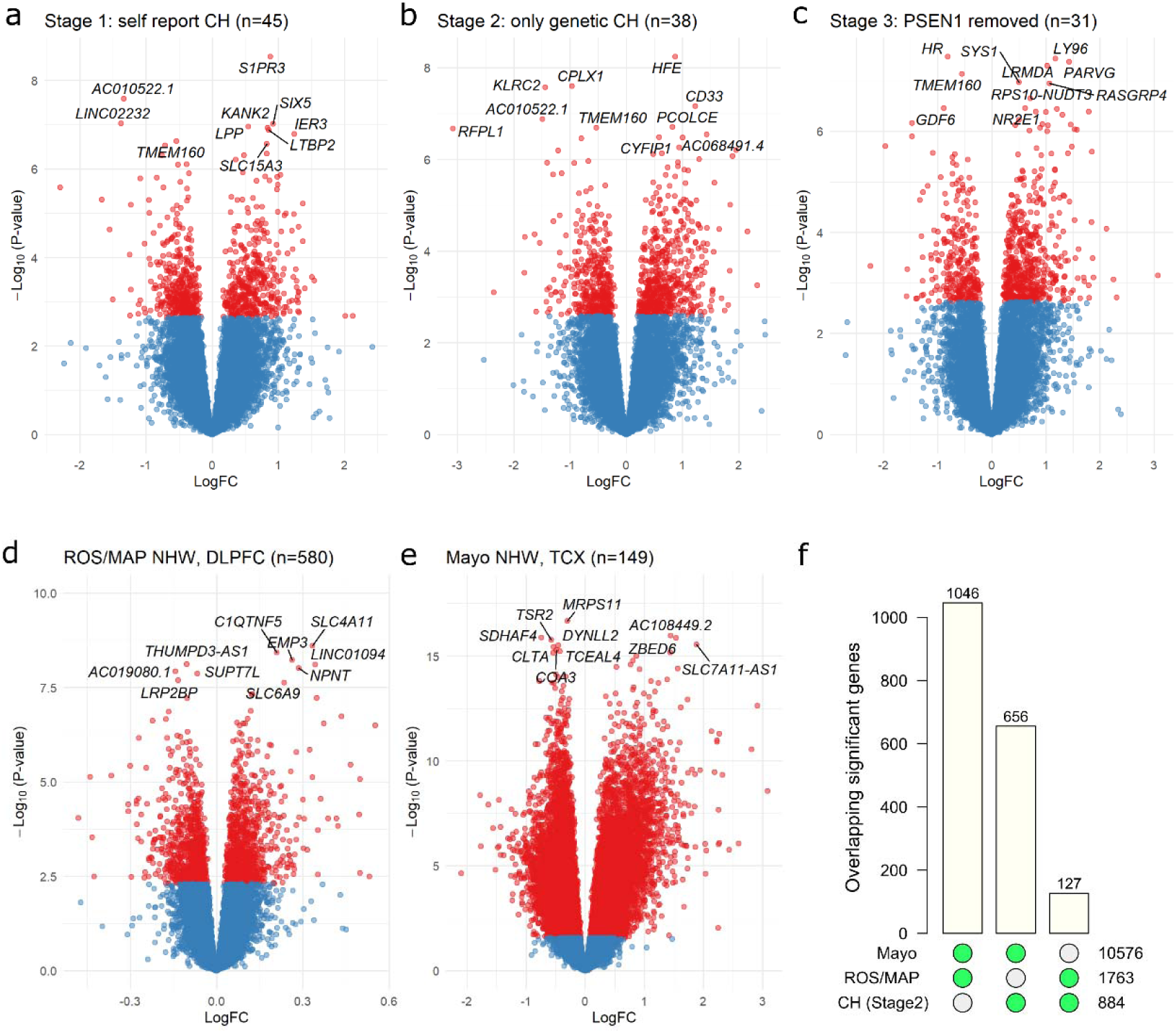
Results of single-gene transcriptome-wide associations with LOAD in Caribbean-Hispanics and non-Hispanic Whites. Gene-level volcano plots for association of genes with LOAD status in A) the full self-reported Caribbean-Hispanic (CH) sample (stage 1, n=45), B) a subset of only genetically-confirmed CH subjects (stage 2, n=38), C) a subset of only genetically-confirmed CH subjects with *PSEN1* mutation carriers excluded (stage 3, n=31), and two independent samples of non-Hispanic whites: D) ROS/MAP and E) the Mayo RNAseq cohort. The top ten significantly differentially expressed genes in each analysis are labeled. The Y-axes indicate two-sided –log_10_(*p*-values) for robust regression testing differential expression The X-axes indicate log_2_ fold-change in expression. F) the number of genome-wide significant differentially expressed genes in common across all three datasets (CH Stage 2 analysis). DLPFC = dorsolateral prefrontal cortex source tissue; LogFC = log_2_ fold-change in expression; TCX = temporal cortex source tissue.

In stage 1, 790/17,300 genes were found to be differentially expressed in LOAD vs. controls at false discovery rate (FDR)-adjusted *p*<0.05 (**Figure 1A**). Stages 2 and 3 found similar numbers of LOAD-associated genes: 884 (out of 17,301) and 817 (out of 17,313), respectively (**Figures 1B-C**). Given that stage 2 found the largest number of significant genes, and that all individuals in stage 2 were genetically-confirmed three-way admixed, we chose to carry forward this model for further analyses (full summary statistics in **Supplementary Table 1**). Transcriptome-wide significant genes encoded several known LOAD loci: *CD33, CD2AP*, and *HLA-DRB1*. At uncorrected *p*<0.05, we identified an additional 11 LOAD loci differentially expressed (*INPP5D, ABI3, FRMD4A, CLU, CR1, FERMT2, TYROBP, FBXL7, MS4A6A, MS4A4A*, and *TREM2 (p=0*.*051*)).

**Table 1.** Sample Demographics for Caribbean-Hispanic and Non-Hispanic White RNAseq Samples.

| Sample | Non-LOAD | LOAD | Diff (*p*) |
| --- | --- | --- | --- |
| **CH** | **(n=16)** | **(n=22)** |  |
| sex (M/F) | 8M, 8F | 6M, 16F | 0.19 |
| age | 66.9 (11.8) | 78.1 (10.2) | 0.0044 |
| RIN | 5.8 (1.5) | 4.2 (1.5) | 0.0018 |
| *PSEN1* G206A status (+/-) | 16- | 7+/15- | 0.014 |
| Library size, millions | 16.4 (4.3) | 15.8 (4.1) | 0.65 |
| **ROS/MAP (NHW)** | **(n=238)** | **(n=358)** |  |
| sex (M/F) | 88M, 140F | 111M, 241F | 0.09 |
| age | 86.9 (7.2) | 90.1 (5.9) | $5.0 \times 10^{-8}$ |
| RIN | 7.2 (1.1) | 7 (0.9) | 0.021 |
| PMI | 7.0 (4.1) | 7.6 (5.2) | 0.11 |
| Library size, millions | 29.7 (9.7) | 26.7 (7) | 0.0024 |
| **Mayo (NHW)** | **(n=69)** | **(n=80)** |  |
| sex (M/F) | 34M, 35F | 31M, 49F | 0.044 |
| age | 82.8 (8.5) | 82.6 (7.7) | 0.9 |
| RIN | 7.7 (1) | 8.6 (0.6) | $1.7 \times 10^{-9}$ |
| PMI | 6.3 (6.5) | 6.6 (4.3) | 0.71 |
| Library size, millions | 41.1 (11.1) | 45.3 (10) | 0.016 |

Note: \*p-values in “Diff” column correspond to those from two-sided, two-sample t-tests for continuous measures (age, RIN, library size, and PMI) and from two-sided Fisher’s exact tests for dichotomous measures (sex and *PSEN1* G206A mutation status). Summary measures for continuous outcomes are means (standard deviations in brackets). RIN = RNA integrity number; LOAD = late-onset Alzheimer’s disease; M = male; F = female; PMI = postmortem interval; CH = Caribbean-Hispanic; NHW = non-Hispanic white; ROS/MAP = Religious Orders Study / Memory and Aging Project; PMI = postmortem interval.

To compare results between CH and NHW populations, we re-analyzed RNA sequencing data from postmortem dorsolateral prefrontal cortex (DLPFC) of 595 NHW subjects from the Religious Orders Study and Memory and Aging Project (ROS/MAP) studies.^5^ A stratified sensitivity analysis was performed to ensure that differences in age range between ROS/MAP and CH samples did not impact our comparisons (**Supplementary Information**). After QC, 580 subjects remained (n_LOAD_=352, n_non-LOAD_=228). We also accessed temporal cortex RNA-sequencing data from the Mayo RNAseq study.^6,7^ After identical QC, 149 subjects remained (n_LOAD_=80, n_non-LOAD_=69). Full report of QC and statistical modelling can be found in **Supplementary Information**. In ROS/MAP and Mayo, differential expression analyses found 1,763/17,665 and 10,576/19,380 significant genes, respectively (**Figure 1D-E;** summary statistics in **Supplementary Tables 2 and 3**). Comparison of these results against published TWAS in these cohorts^8–10^ (**Supplementary Information; Supplementary Figure 2 and 3**) validated these findings. Thus, we carried forward results from three independent RNAseq datasets, including two different brain regions and two different ethnicities (see **Table 1** for sample demographics).

**Table 2.**
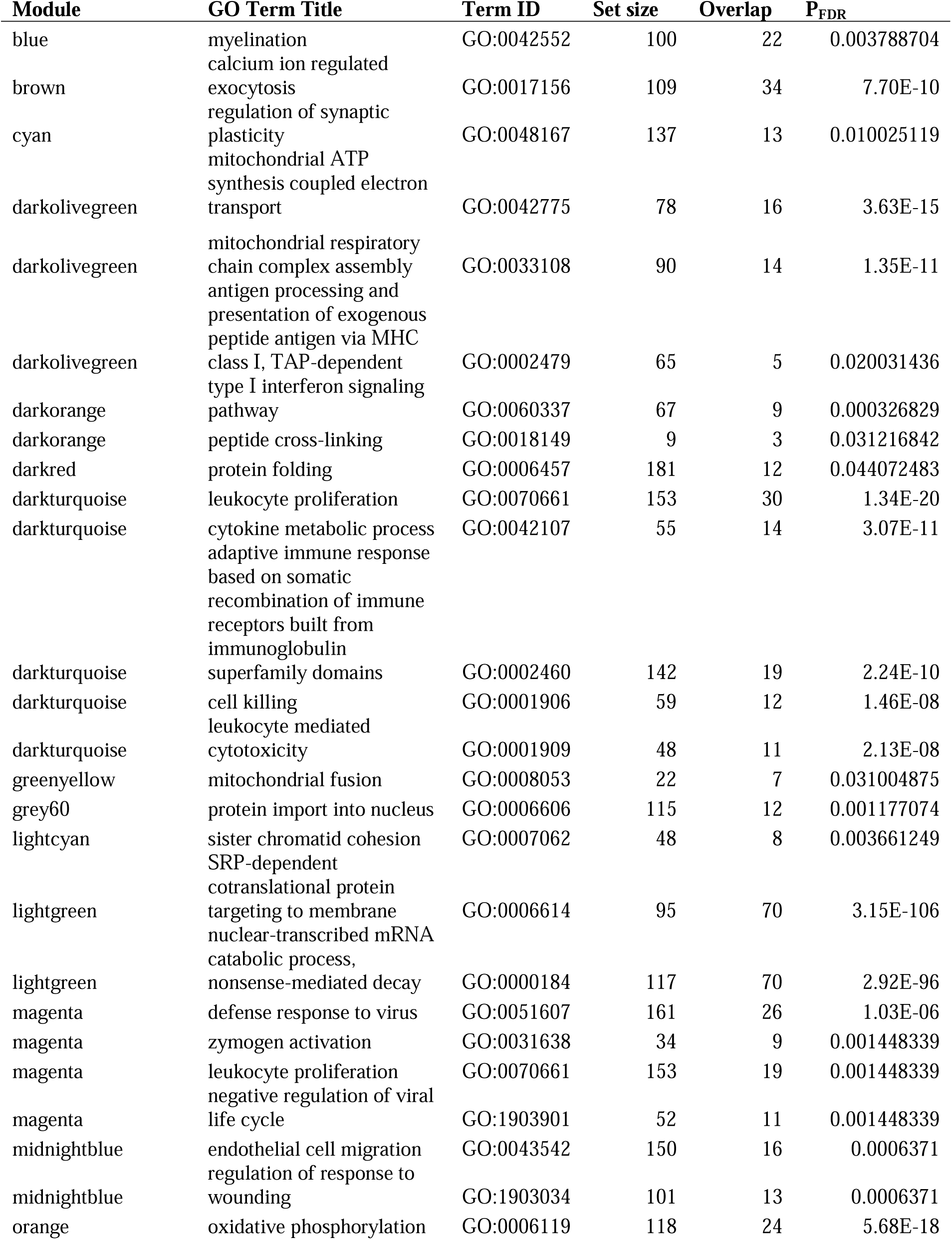
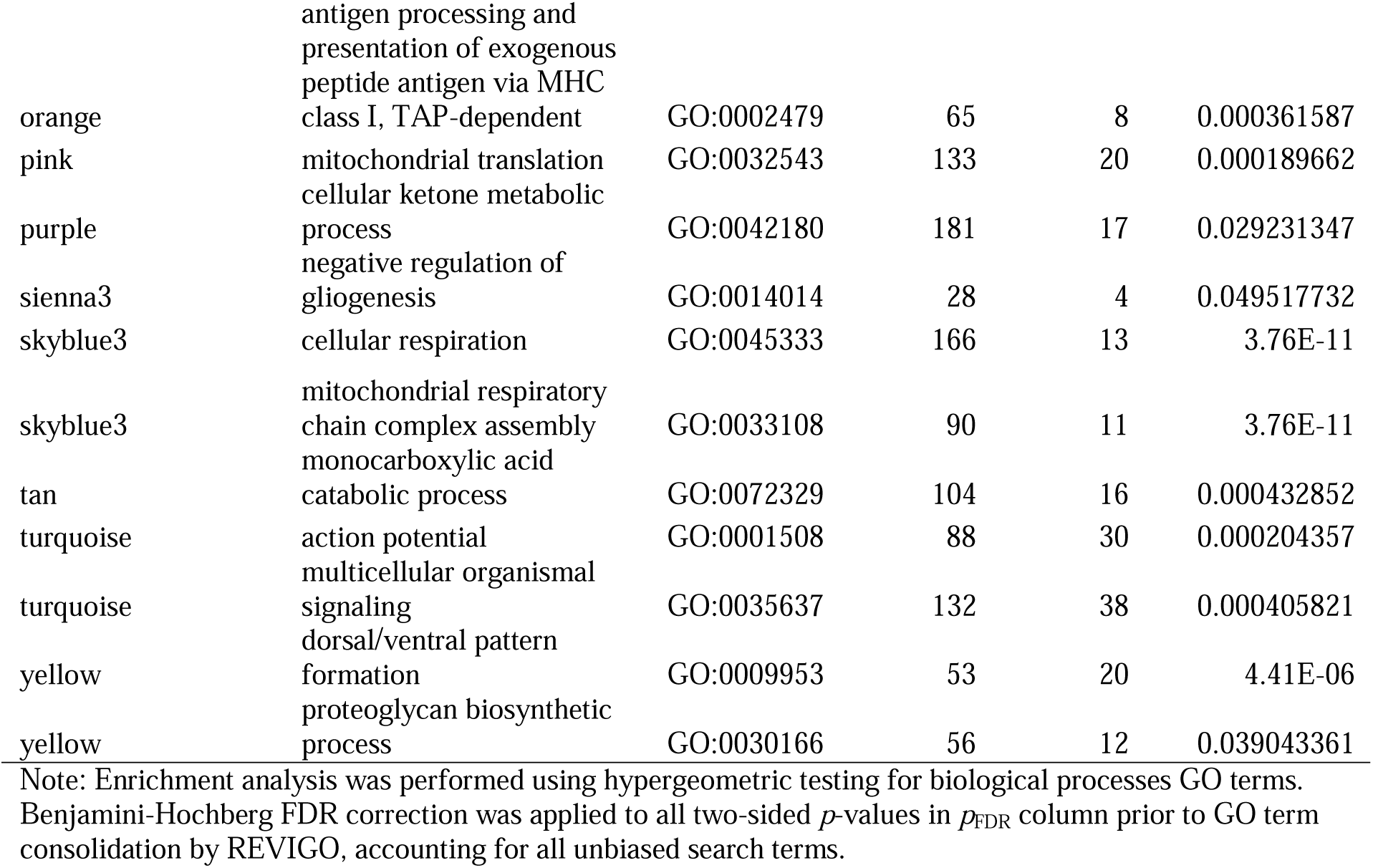
Consensus gene co-expression module GO enrichment consolidated results for biological processes.

**Figure 2.**
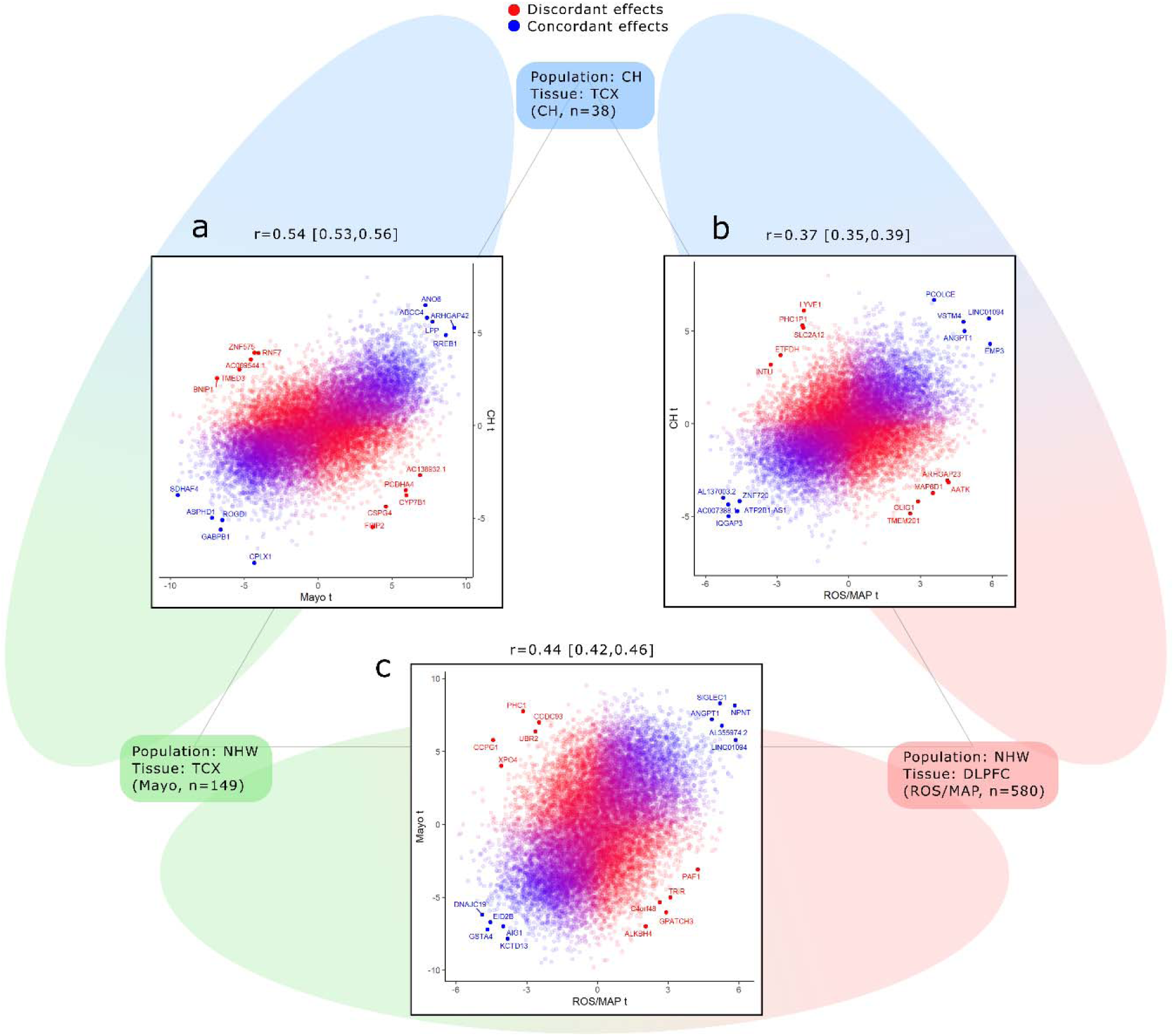
Correlation of gene-wise differential expression effects between each study. A) Scatterplot showing the correlation between moderated t-statistics from robust regression of differential expression in the CH (Y-axis) and Mayo cohort (X-axis) studies. B) Scatterplot showing the same comparison for CH (Y-axis) and ROS/MAP (X-axis), and C) the same comparison for the Mayo cohort (Y-axis) and ROS/MAP (X-axis). Points are colored according to the rank of concordance or discordance of their effects (calculated at the product of t-statistics between the two studies being compared). The top 5 concordant and discordant genes in each direction are labeled in each plot. Above each plot are the Pearson correlation coefficient for moderated t-statistics and its 99% confidence interval. DLPFC = dorsolateral prefrontal cortex source tissue; ROS/MAP = Religious Orders Study and Memory and Aging Project; TCX = temporal cortex source tissue.

**Figure 3.**
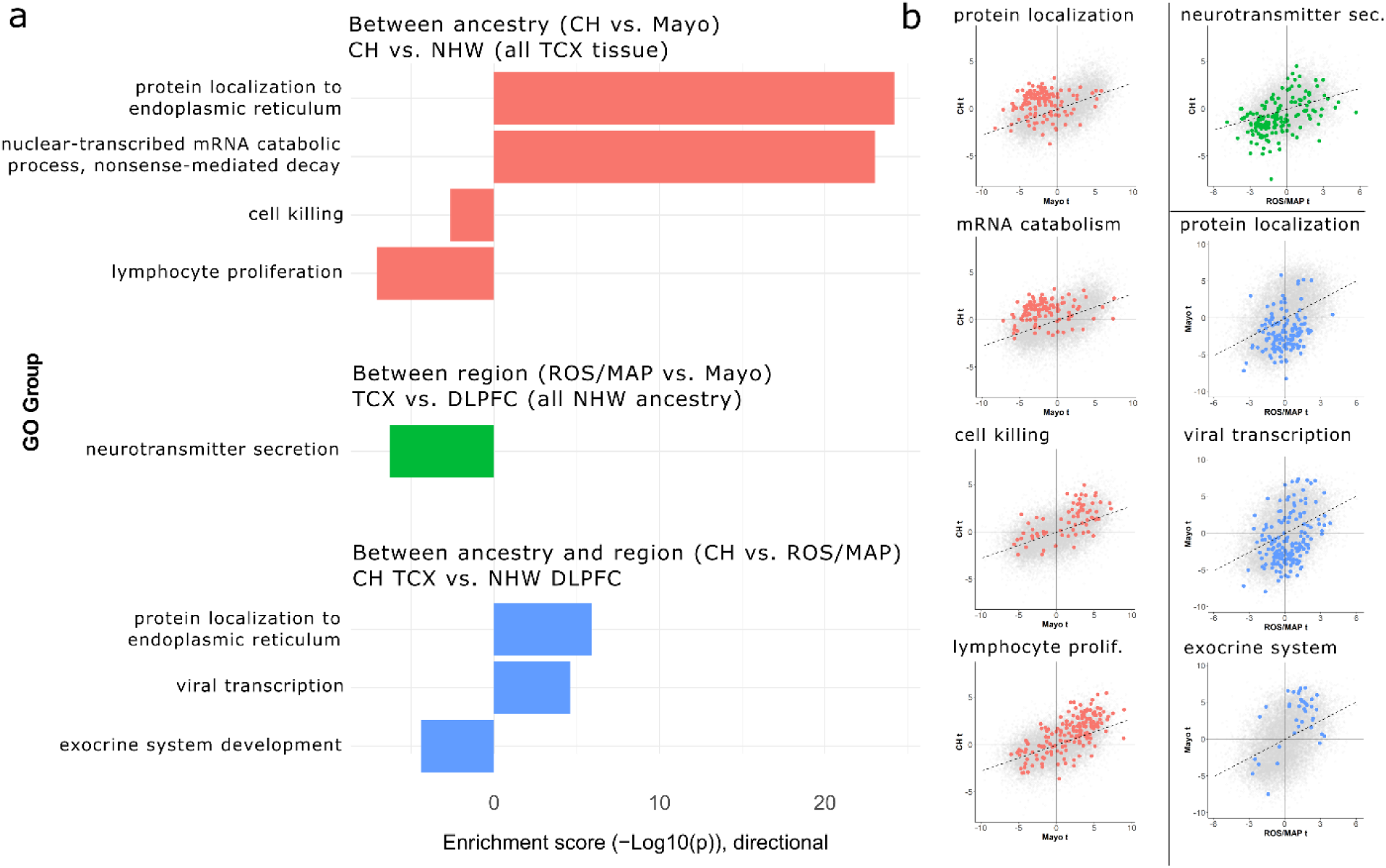
Enrichment analyses for genes with concordant and discordant LOAD effects between ancestry and brain region tissue source. A) Barplot summarizing AUC-based GO enrichment analysis using tprod ranks for all FDR-significant GO terms with a REVIGO dispensability score of 0. The X-Axes indicate the enrichment –log10(*p*-values), with values above 0 indicating enrichment toward higher ranks (greater discordance of between-sample effect) and values below 0 indicating enrichment toward lower tanks (greater concordance of between-sample effect). B) Scatterplots for each enrichment in panel A, showing the LOAD differential expression t-values for genes belonging to each enriched gene set within the context of all genes. Genes belonging to the labeled GO set are colored to match the barplot in A. CH = Caribbean-Hispanic; FDR = false discovery rate; GO = gene ontology; DLPFC = dorsolateral prefrontal cortex source tissue; ROS/MAP = Religious Orders Study and Memory and Aging Project; TCX = temporal cortex source tissue.

We initially focused on genes with ancestry-independent effects, i.e. replicated across Caribbean Hispanic and at least one non-Hispanic white cohort. Considering the number of significant genes per dataset, we observed a greater proportion of replicated signals between Caribbean Hispanic and Mayo (656 in common out of 884 in CH = 74% replication) than between CH and ROS/MAP (127/884 = 14% replication), suggesting that differences in brain region supersede those due to ancestry. In total, 114 genes were genome-wide significant in all three TWAS; 111 with concordant directions of effect. Among these 111 concordant genes, several were of known significance to LOAD, including *LINC1904, FYN, VIM, NFKB1, LTBP2, KHDRBS1, PLCE1, CRK*, and *SYNM*. At uncorrected *p*<0.05, 863 genes were significant in all three TWAS, 808 with concordant directions of effect. Notably, we found the known CH GWAS-implicated LOAD risk gene *FBXL7*^11^ overexpressed in LOAD at uncorrected *p*<0.05 in all three samples. In addition, *CD33* was the second most strongly LOAD-overexpressed gene in CH (*p*_FDR_=3.0×10^−4^) as well as genome-wide significant in Mayo (*p*_FDR_=3.1×10^−3^).

We then aimed to identify genes with ancestry-specific effects by combining summary statistics from genes common across the three studies (n_genes_=15,857) and comparing effects between 1) CH vs. ROS/MAP, 2) CH vs. Mayo, 3) ROS/MAP vs. Mayo (**Figure 2**). We then performed AUC-based gene ontology (GO) term enrichment, with resulting significant GO groups consolidated to minimize redundancy and overlap using the REVIGO tool.^12^ Pairwise comparisons of moderated t-statistics revealed moderate positive correlations (CH vs. Mayo r=0.54, C.I._99%_=[0.53,0.56]; CH vs. ROS/MAP r=0.37, C.I._99%_=[0.35,0.39]; ROS/MAP vs. Mayo (r=0.44, C.I._99%_=[0.42,0.46]), suggesting again that sharing of LOAD-related molecular changes are more pronounced when comparing the same brain region. We then ranked genes according to concordance of effect between samples, specifically, by the product of their t-statistics (“tprod”) for association with LOAD, assigning the highest ranks to genes with the largest discrepancy in direction and effect size between NHW and CH. Thus, higher ranks indicate sample-specific effects vs. lower ranks indicating common effects.

Enrichment analyses revealed eight unique, non-redundant (REVIGO dispensability = 0) biological processes with either significantly discordant or concordant LOAD-related effects (**Figure 3)**. First, comparing CH vs. Mayo, those genes ranking highly were significantly enriched for “protein localization to endoplasmic reticulum” (*p*_FDR_=6.5×10^−25^) and “nuclear−transcribed mRNA catabolic process, nonsense−mediated decay” (*p*_FDR_=9.3×10^−24^), whereas low-ranking genes were enriched for “lymphocyte proliferation” (*p*_FDR_=8.2×10^−8^) and “cell killing” (*p*_FDR_=2.4×10^−3^). Notably, the “response to beta-amyloid” (*p*_FDR_=1.4×10^−4^) category was also concordantly downregulated in both datasets, though it was not assigned a dispensability score of 0 (disp=0.057). Second, ROS/MAP vs. Mayo, only “neurotransmitter secretion” (*p*_FDR_=8.2×10^−8^) was significantly enriched with concordant downregulation. Finally, comparing CH vs. ROS/MAP, “protein localization to endoplasmic reticulum” (*p*_FDR_=8.2×10^−8^) and “viral transcription” (*p*_FDR_=8.2×10^−8^) both showed significant discordance, whereas “exocrine system development” (*p*_FDR_=8.2×10^−8^) was concordant. Importantly, three of the eight GO groups identified had significant overlap in gene membership, namely “protein localization to endoplasmic reticulum”, “nuclear−transcribed mRNA catabolic process, nonsense−mediated decay “, and “viral transcription”, with 78 genes in common (**Supplementary Figure 4**). These 78 genes belong to the class of large and small ribosomal subunits, with the strongest representation in the “SRP-dependent co-translational protein targeting to membrane” GO ontology (78/98 genes).

**Figure 4.**
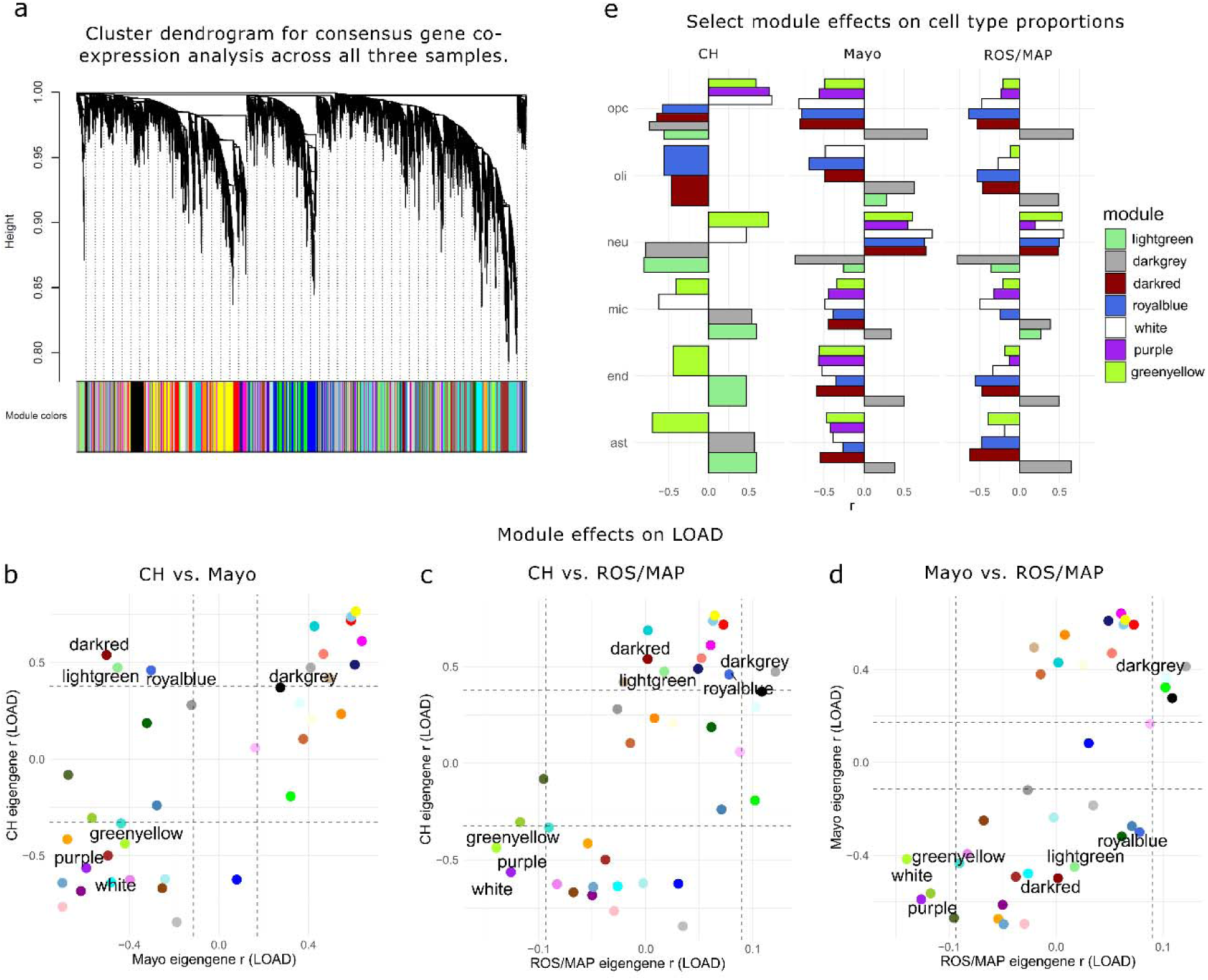
Consensus gene co-expression module analysis with LOAD and cell type proportions. A) Clustered dendrogram showing the common gene set hierarchical structure and consensus gene module definitions. Panels B-D show consensus module eigengene associations with LOAD (Pearson r) for all modules compared between pairwise sample combinations. Modules with concordant effects (*p*_FDR_<0.05 with same direction of effect) in all three samples, or significantly discordant (*p*_FDR_<0.05 in at least two samples but with opposite direction of effect), are labeled; four modules were significantly associated with LOAD in all three samples; greenyellow, purple, and white modules were consistently downregulated, while the darkgrey module was upregulated. In contrast, three modules showed significant but directionally discordant effects on LOAD, all when comparing CH to Mayo; darkred, royalblue, and lightgreen modules were all significantly upregulated in LOAD in CH and downregulated in Mayo. Colors correspond to module definitions. F) Association (Pearson r) of discordant and concordant module eigengenes with estimated cell type proportions from BRETIGEA in each sample. Only associations significant after FDR correction are shown. CH = Caribbean-Hispanic; ROS/MAP = Religious Orders Study and Memory and Aging Project; ast = astrocytes; end = endothelial cells; mic = microglia; neu = neurons; oli = oligodendrocytes; opc = oligodentrocyte precursor cells.

Finally, we sought to characterize network-level effects of expression between ancestry and brain region. First, consensus gene co-expression modules were defined across all three samples using Weighted Gene Co-Expression Analysis (WGCNA).^13^ Then, modules were characterized for GO enrichment and tested for association with LOAD and cell type proportions in each sample separately to identify sample-specific processes. Cell type proportions were estimated using human marker genes from the Brain Cell Type Specific Gene Expression Analysis (BRETIGEA)^14^ package (see **Supplementary Information**). A total of 39 discrete consensus co-expressed gene modules were identified, ranging from 43 to 1,710 genes in size (**Figure 4A; Supplementary Table 4)**, with 21 modules significantly enriched for at least one biological process (**Table 2;** extended enrichment results for uncorrected *p*<0.05 in **Supplementary Table 5**). Association tests of module eigengenes with LOAD in each dataset revealed largely conserved network-level effects: comparing CH and Mayo, LOAD effects were most strongly correlated (r=0.72, *p*=2.7×10^−7^), followed by Mayo vs. ROS/MAP (r=0.67, *p*=3.7×10^−6^), and CH vs. ROS/MAP (r=0.60, *p*=6.3×10^−5^). Several modules with concordant and discordant LOAD associations were identified (see **Figure 4B-E;** full association statistics in **Supplementary Table 6**), among which the lightgreen module, highly enriched for “signal-recognition particle (SRP)-dependent cotranslational protein targeting to membrane” (*p*=3.2×10^− 106^), was positively associated with endothelial, microglial, and astrocytic cells in CH, but not in ROS/MAP or Mayo, suggesting an ancestry-specific molecular signature of these cell types. To our knowledge, this is the first investigation of gene expression from postmortem LOAD brain in CH individuals. Our results show a substantial overlap in LOAD-related genes between CH and NHW, particularly in pathways related to immune cell proliferation, cell killing, and neurotransmitter secretion, whereas differences highlight roles of ribosomal genes and those involved in viral transcription. We speculate that (SRP)-dependent protein targeting may be disproportionately perturbed in CH LOAD patients given the strong ethnic-specific enrichment at both the gene and module. Intriguingly, SRP-dependent genes are strongly dysregulated periodontitis affected periodontal tissue.^15^ The main driver of periodontitis pathogenesis, P. gingivalis, is known to secrete gingipains which have been linked to LOAD pathogenesis in mice^16^ and humans.^17^ Understanding ancestry-specific molecular profiles of LOAD brain tissue are a first step toward developing research questions and eventually interventions effective in non-Caucasian high risk populations.

Key limitations in this study include different methods of tissue ascertainment and study design that ultimately impact the results; ROS and MAP are community-based, prospective cohort studies, whereas the CH and Mayo samples are case-control designs selected for diagnosis. Additionally, samples were not scanned for rare LOAD mutations using whole genome sequencing and CH specifically was a relatively small sample collected over several decades. The difficulties of ascertaining brains in specific populations challenge the availability of large samples sizes and tissue collected over a short period of time (i.e. higher quality). This resulted in samples with relatively low RNA quality (median RIN=4.5). Hispanics tend to not participate in either organ donation in general, or brain donation more specifically, to the same extent as non-Hispanic Whites (NHW).^18^ Nevertheless, by performing ribosomal RNA depletion prior to sequencing, we ensured that the impact of low RNA quality was mitigated. In fact, ribosomal RNA depletion has been shown to perform very well^19,20^ at amounts far below recommendation and over a wide range of intact and degraded material. Despite these challenges, our analyses proved reliable by showing a moderate concordance between results from all three samples at the genome-wide scale. We were also able to identify several well-known LOAD-associated loci from previous GWAS and sequencing studies. Importantly, *FBXL7* was previously identified by a GWAS from our group.^11^ We also found an increased expression of *CD33* in CH LOAD brains, consistent with the effect of the CD33 risk allele which increases the level of expression of full length CD33 in myeloid cells.^21^

In sum, we performed RNA-sequencing on postmortem brain from a small sample of Caribbean-Hispanic elderly and two large independent non-Hispanic Whites cohorts, ultimately identifying candidate genes and processes that are consistently dysregulated across ethnicity or show ethnic-specific effects. Further work in large admixed cohorts will permit a deeper understanding of ancestry-specific mechanisms that may be used to predict risk, onset of pathology, and potentially provide precision treatment options.

## Acknowledgements

This work was supported by grants from the National Institutes of Health: R21AG054832, R56AG059756, R01AG056531, P50AG008702, P30AG10161, R01AG15819, R01AG17917, U01AG61356, R01AG015473, R56 AG063908, P30AG066462, R01NS095922 and P50AG0008702. Funding support for DF was provided by The Koerner Family Foundation New Scientist Program.

## Data Availability

RNA sequencing and LOAD phenotype data for the ROS/MAP and Mayo cohorts are available via approved access at the Synapse AMP-AD Knowledge Portal (https://adknowledgeportal.synapse.org/, doi: 10.7303/syn2580853).

## Author Contributions

DF was responsible for data curation from new and published sources, data quality control and analyses, manuscript writing and editing. SS contributed to data preprocessing, interpretation, and manuscript editing. LF contributed to several aspects of the enrichment analysis, study design, and drafting of the manuscript. ISM was responsible for preparation of CH samples for RNA sequencing, quality control, and manuscript editing. JAS and DAB were responsible for ROS/MAP postmortem data collection, PLDJ was responsible for ROS/MAP RNA sequencing data generation, and all three authors contributed to elements of study design and manuscript editing. RM and GT were responsible for CH postmortem data ascertainment, contributed to study design, analytics design, and drafting of the manuscript.

