## Supplementary file for "The Caribbean-Hispanic Alzheimer’s Brain Transcriptome Reveals Ancestry-Specific Disease Mechanisms"

### Description of Supplementary Files

#### Supplementary Table 1

Full summary statistics for CH sample LOAD differential expression analysis. logFC=log_2_ fold-change for differential expression; AveExpr=mean sample expression (log2(CPM)); t=moderated t-statistic for LOAD association; P.Value=uncorrected association *p*-value (two-sided); adj.P.Val=FDR-corrected association *p*-value; se=standard error of LogFC; gene=Ensemble gene ID; hugo=HUGO Gene Nomenclature Committee (HGNC) gene symbol; chr=gene chromosome.

#### Supplementary Table 2

Full summary statistics for ROS/MAP sample LOAD differential expression analysis. Column definitions identical to Supplementary Table 1.

#### Supplementary Table 3

Full summary statistics for Mayo sample LOAD differential expression analysis. Column definitions identical to Supplementary Table 1.

#### Supplementary Table 4

Definitions of WGCNA consensus modules derived across CH, ROS/MAP, and Mayo. Module=module label; gene= Ensemble gene ID; hugo= Nomenclature Committee (HGNC) gene symbol.

#### Supplementary Table 5

Results of WGCNA consensus module GO enrichment analysis for all module-GO term (biological processes) enrichment reaching uncorrected *p*<0.05. Title=GO term full title; overlap=number of genes in module and GO group; setSize= number of genes in GO group; hitListSize=number of genes in module; P.Value=uncorrected enrichment *p*-value (two-sided); adj.P.Val=FDR-adjusted enrichment p-value; ID=GO ID; rank=within-module rank; module=consensus module label.

#### Supplementary Table 6

Results of module-trait associations in CH, ROS/MAP, and Mayo cohorts. Module=module label; r=Pearson correlation coefficient; variable=outcome trait (BRETIGEA cell type estimates and LOAD status); p=uncorrected *p*-value (two-sided); fdr_within=within-sample FDR-corrected *p*-values; sample=study sample; fdr_all=overall across-study FDR-corrected *p*-value.

### Supplementary Methods

#### 1. Study subjects, sample preparation, and RNA sequencing

##### 1.1. Sample 1: Caribbean-Hispanics (CH)

The CH sample was ascertained from the brain bank of the Alzheimer's Disease Research Center (ADRC) at Columbia University (New York, NY, USA). All ADRC participants are informed of the opportunity to participate in the brain bank. This research clinic referral-based brain bank consists of over 600 brains from autopsies performed between 1989–2016. The anatomical origin of each sample is recorded using the map of Brodmann for cortical samples. The samples are frozen using liquid nitrogen vapor (LNV) at −160°C, and barcode labeled; three measurable parameters serve to assess the quality of the tissue banked: the pH value, the yield of total RNA per unit of tissue, and the extent of degradation of ribosomal RNA. LNV fresh frozen blocks are available for most of the cases (size range between 0.3 × 0.5 × 1.0 cm and 0.5 × 2.5 × 3.0 cm), which are especially suitable for studies requiring preservation of the cellular morphology and that of the cytoarchitecture of the area of interest. Also available are aliquots of fresh-frozen pulverized brain parenchyma (1.0–1.5 ml), especially for studies focusing on biochemistry, protein studies, or molecular biology. The latter were requested for this study. In total, 45 cases from Temporal cortex (TCX) were collected. Cases were selected if age>50 years old, and neuropathologically defined LOAD or control without any neuropathological diagnosis (see below) and reported Hispanic ancestry. The ADRC protocol was approved by the institutional review boards of the New York State Psychiatric Institute and Columbia University.

###### Diagnosis of LOAD

Details of the neuropathological assessment have been extensively described elsewhere.^1,2^ Neuritic plaques and neurofibrillary tangles were assessed using hematoxylin-eosin and the modified Bielschowsky silver method. Immunohistochemistry was performed for beta amyloid, phosphorylated tau, ubiquitin (p62), TDP-43 and glial fibrillary acidic protein. Briefly, all LOAD cases met the criteria of Braak stage ≥4 neurofibrillary pathological findings and also met the CERAD neuropathological criteria for definite LOAD.

###### Sample preparation and RNA sequencing

Total RNA was extracted after bead based homogenization and using Qiagen RNeasy Plus Mini Kit (Qiagen). RNA samples were quantified using Qubit 2.0 Fluorometer (Life Technologies, Carlsbad, CA, USA) and RNA integrity was checked with 4200 TapeStation (Agilent Technologies, Palo Alto, CA, USA). RNA library preparations, sequencing reactions, and bioinformatics analysis were conducted at GENEWIZ, LLC. (South Plainfield, NJ, USA).

rRNA depletion was performed using Ribozero rRNA Removal Kit (Human/Mouse/Rat probe) (Illumina, San Diego, CA, USA). RNA sequencing library preparation used NEBNext Ultra RNA Library Prep Kit for Illumina by following the manufacturer’s recommendations (NEB, Ipswich, MA, USA). Briefly, enriched RNAs were fragmented for 15 minutes at 94 °C. First strand and second strand cDNA were subsequently synthesized. cDNA fragments were end repaired and adenylated at 3’ends, and universal adapter was ligated to cDNA fragments, followed by index addition and library enrichment with limited cycle PCR. Sequencing libraries were validated on the Agilent TapeStation (Agilent Technologies, Palo Alto, CA, USA), and quantified using Qubit 2.0 Fluorometer (Invitrogen, Carlsbad, CA) as well as by quantitative PCR (Applied Biosystems, Carlsbad, CA, USA).

The sequencing libraries were clustered on six lanes of a flowcell. After clustering, the flowcell was loaded on the Illumina HiSeq instrument according to manufacturer’s instructions. The samples were sequenced using a 2x150 Paired End (PE) configuration. Image analysis and base calling were conducted by the HiSeq Control Software (HCS). Raw sequence data (.bcl files) generated from Illumina HiSeq were converted into fastq files and de-multiplexed using Illumina's bcl2fastq 2.17 software. One mismatch was allowed for index sequence identification.

###### CH genotyping and global admixture estimation.

CH individuals were genotyped employing the Infinium Global Screening Array-24 Kit (<https://www.illumina.com/products/by-type/microarray-kits/infinium-global-screening.html>). Imputation was performed using a validated pipeline for admixed populations; individuals labeled as mutation carriers in the Brain Bank records were confirmed by imputing the *PSEN1* G206A variant as described in detail elsewhere.^3^ It has to be noted that the CH sample has not received whole-genome or whole-exome sequencing, therefore we could not identify exhaustively the entire pool of known mutations or large-effect sizes risk variants (e.g. *TREM2*). This is a caveat held in common with most other published work of RNAseq datasets, including analyses of ROS/MAP and Mayo cohort data.

Global admixture was estimated for CH individuals using the ADMIXTURE (v1.3.0) software. We conducted supervised admixture analyses using African Yoruba (“YRI”), European (“EUR”) and Native American subjects (“NAT” - eight Surui, 21 Maya, 14 Karitiana, 14 Pima and seven Colombian) from the Human Genome Diversity Project (HGDP) as surrogates for European, African and Native American ancestry, respectively. We used ~100,000 autosomal SNPs that were I) common (i.e. MAF >1 %) and II) in linkage equilibrium. We retained individuals that showed at least 1% of each ancestral component (“three-way admixed” - **Supplementary Figure 1**).

##### 1.2. Sample 2: Religious Orders Study and Memory and Aging Project Non-Hispanic Whites (NHW; ROS/MAP)

The sample characteristics of the ROS/MAP cohort subset studied here have been published in detail elsewhere.^4^ Briefly, ROS and MAP are two longitudinal cohort studies of the elderly, one recruiting from regions across the United States and the other only from the greater Chicago area. All subjects are recruited without signs of dementia (mean age at entry = 78 ± 8.7 (SD) years) and agree to annual clinical and neurocognitive evaluation. In addition, all participants sign an Anatomical Gift Act allowing for brain autopsy at time of death, which permits multi-omic investigations of bulk brain tissue from multiple regions. All study participants provided informed consent and both studies were approved by the Rush University Institutional Review Board.

###### Diagnosis of LOAD

For defining LOAD diagnosis, details have been published previously.^5^ Briefly, brains were removed and examined by a board-certified neuropathologist blinded to clinical data. Brains were cut into 1cm coronal slabs and fixed for at least 3 days in 4% paraformaldehyde. We used defined landmarks to obtain tissue from dorsolateral prefrontal cortex, middle and inferior temporal cortex, inferior parietal, hippocampus CA1/subiculum, entorhinal cortex proper, ventromedial caudate, and posterior putamen. Tissue blocks were processed, paraffin embedded, cut into sections, and mounted on glass slides. Neuropathologic diagnoses were made by a board-certified neuropathologist blinded to age and clinical data. Bielschowsky silver stain 6 micron sections were used to visualize neuritic plaques, diffuse plaques, and neurofibrillary tanglesin the frontal, temporal, parietal, entorhinal, and hippocampal cortices, as previously described.^6^ A neuropathologic diagnosis of “no LOAD,” “low likelihood LOAD,” “intermediate likelihood LOAD,” or “high likelihood LOAD” was given based on semiquantitative estimates of neuritic plaque density as recommended by CERAD and Braak staging as recommended by the National Institute on Aging (NIA)-Reagan criteria. For analyses, a neuropathologic diagnosis of LOAD was assigned if NIA-Reagan diagnosis was either intermediate or high likelihood.

###### Sample preparation and RNA sequencing

For mRNA quantification, library construction and sequencing have been described in detail previously.^7^ Briefly, tissue samples from DLPFC were analyzed by the Broad Institute’s Genomics Platform for transcriptome library construction following the dUTP protocol^8^ and Illumina sequencing. Five micrograms of total RNA as measured by RiboGreen at a concentration of 50 nanogram/microliter with RNA Integrity Number (RIN) score of 5 or better were submitted for cDNA library construction. In total, 726 subjects were sequenced across 10 batches using both the dUTP method, barcoded and pooled for sequencing, and the newer Illumina TruSeq method modified by The Broad Institute Genomics Platform to be strand specific and to use larger insert sizes (which results in libraries closely resembling those obtained by the dUTP method). The Truseq method uses only 250 nanograms of RNA input. Sequencing was carried out using the Illumina HiSeq2000 with 101 bp paired end reads for a targeted coverage of 50M paired reads. All subsequent analyses controlled for batch to mitigate methodological biases.

##### 1.3. Sample 3: Mayo RNAseq Study Non-Hispanic Whites (NHW; Mayo)

The Mayo RNAseq cohort has been described in detail.^9,10^ In total, 276 Temporal cortex (TCX) samples were collected from 312 North American Caucasian subjects with neuropathological diagnosis of AD, progressive supranuclear palsy (PSP), pathologic aging (PA) or elderly controls (CON) without neurodegenerative diseases. Within this cohort, all LOAD subjects were from the Mayo Clinic Brain Bank (MCBB). Thirty-one control TCX samples were from the MCBB, and the remaining control tissue was from the Banner Sun Health Research Institute.

###### Diagnosis of LOAD

All subjects selected from the MCBB and Banner underwent neuropathologic evaluation by Dr. Dennis Dickson or Dr. Thomas Beach, respectively. All LOAD participants had definite diagnosis according to the NINCDS-ADRDA criteria and had Braak neurofibrillary tangle (NFT) stage of IV or greater. Control subjects had Braak NFT stage of III or less, CERAD neuritic and cortical plaque densities of 0 (none) or 1 (sparse) and lacked any of the following pathologic diagnoses: LOAD, Parkinson’s disease, dementia due to Lewy bodies, vascular dementia, PSP, motor neuron disease, corticobasal degeneration, Pick’s disease, Huntington’s disease, fronto-temporal lobar degeneration, hippocampal sclerosis or dementia lacking distinctive histology.

###### Sample Preparation and RNA Sequencing

For sequencing, Mayo Clinic RNAseq samples were assigned to flowcells randomized for age at death, sex, RIN, Braak stage and diagnosis. Library preparation and sequencing of the samples were conducted at the Mayo Clinic Medical Genome Facility Gene Expression and Sequencing Cores, as previously described.^9^ The TruSeq RNA Sample Prep Kit (Illumina, San Diego, CA) was used for library preparation from all samples. The library concentration and size distribution was determined on an Agilent Bioanalyzer DNA 1000 chip. Three samples were run per flowcell lane using barcoding. All samples underwent 101 base-pair (bp), paired-end sequencing on Illumina HiSeq2000 instruments. The Mayo cohort research team has performed their own quality control of samples available on the Synapse sharing hub, including flags for sex mismatches, cryptic relatedness to other samples, and failing to meet control criteria due to Braak staging. Prior to count-based quality control described below, subjects flagged by Mayo QC were excluded from the sample (except for three subjects who were excluded as PCA outliers, since we performed our own PCA QC procedure on the case-control subset analyzed here).

#### 2. Sequencing alignment and quality control

Raw sequence data were processed identically for CH and ROS/MAP samples. FASTQ files from sequencing experiments were processed batch-wise using the same operational pipeline using FastQC (v0.11.5)^11^, Picard tools (v2.17.4)^12^, STAR aligner (v2.5.3a)^13^, and RSEM (v1.2.31)^14^ against the GRCh38.91 reference genome. For Mayo, base-calling was performed using Illumina’s RTA 1.17.21.3. FASTQ sequence reads were aligned to the human reference genome using TopHat 2.0.12^15^ and Bowtie 1.1.0,^16^ and Subread 1.4.4 was used for gene counting.^17^ FastQC was used for quality control (QC) of raw sequence reads, and RSeQC was used for QC of mapped reads.

#### 3. Post-processing of expected gene counts and quality control for differential expression

While harmonized summary statistics for differential expression in LOAD have been generated for both ROS/MAP and the Mayo cohort as part of the AMP-AD reprocessing initiative^18^, we chose to use raw expression count values for the Mayo cohort to ensure consistency in our statistical approach across all three samples (we also performed comparison of our CH results with the AMP-AD reprocessing statistics, described below in “Validation of new TWAS results against existing analyses in ROS/MAP and Mayo cohorts”).

Gene counts for all three studies were post-processed identically. Counts were read into R (v3.6.3) for processing with edgeR^19^ and limma/voom.^20,21^ First, any genes with a median expected count lower or equal to 15 were excluded. Second, principal components analyses were performed on each batch separately, and subjects flagged for pruning if deviating from the sample median of any of the first 5 principal components by more than 3 interquartile ranges. For the CH sample, PCA was performed on all subjects together, rather than by individual batch, due to small batch size (batch #2 n=5). Notably, known *PSEN1* mutation carriers did not deviate from sample medians in any of these components. Third, gene expected counts were log2 transformed with an offset of 0.5 and each gene evaluated for extreme outlying observations separately. For each gene, any log2(expected count) value deviating from the sample median by 3 interquartile ranges was coerced to its nearest most extreme value within that range. For the much smaller CH sample, this threshold was slightly relaxed to 4 interquartile ranges. Log2(expected counts) were then transformed back to the expected count scale and returned to their DGE object format for the calculation of TMM normalization values (using edgeR calcNormFactors) and mean-variance derived observational-level weights for linear modelling using limma/voom.

#### 4. Robust linear modeling of expression data

The limma package ‘lmFit’ function was used to model log2(expected counts) as a linear function of pathological LOAD status, sequencing batch (brain bank source and flowcell included for Mayo), age, sex, RIN, postmortem interval (not available for the CH sample), percent of usable bases, percent duplicated reads, median 3’ bias, and percent of mapped ribosomal bases. In CH models that included *PSEN1* mutation carriers, a parameter was added to model its effect. For the Mayo cohort, age at death for those 90 or older were only made available as a single level (“90_or_above”) to avoid inadvertent identification of the most elderly subjects; all subjects in this category were assigned a numeric age at death of 90. Further, for Mayo, 42 subjects were missing values for PMI, so, to harness the available data, missing values for this co-variate were mean-imputed prior to modeling. Robust linear modeling was used, allowing for a high number (10,000) of iterations to reach convergence. Significance of the contrast between AD and non-AD status (AD vs. Control for Mayo) was performed using empirical Bayes moderation (eBayes function). In the relatively small CH sample, a modest degree of multi-collinearity (assessed as variance inflation factors exceeding 3) among co-variates was noted in our models of gene expression at the module and individual gene level – the consequences of this correlation structure were explored using partial least squares regression, which determined that the estimates of differential expression presented here are robust, as effect sizes were strongly correlated between both analyses (r>0.95; data not shown).

#### 5. Sensitivity analysis for age differences between ROS/MAP and CH

Due to known transcriptomic differences due to age, and the existence of a substantial number of ROS/MAP participants with age at death greater than the maximum observed in the CH sample (CH age range=50-93; ROS/MAP age range=67-106), we performed a second ROS/MAP differential expression analysis for LOAD in the same manner as described above excluding subjects with age at death greater than 93 (n=459), again with the same co-variates (including age at death). In this analysis, a total of 1,212 genes were significant after FDR correction, 1,077 of which (89%) were also significant in the original, unpruned analysis of all 580 subjects. Moderated t-statistics for the two analyses were very strongly correlated with Pearson r=0.962, C.I._99%_=[0.961,0.964]. Compared to CH, these age-limited ROS/MAP sample results had similar albeit smaller overlap of FDR-significant genes, with 92 (10%) vs. 127 (14%) genes replicating from CH to ROS/MAP, and the correlation of LOAD effects between the age-limited sample results and the CH cohort results was unchanged (CH vs. ROS/MAP r=0.37, C.I._99%_=[0.35,0.39]). Importantly, LOAD-related effects of the *FBXL7* gene were preserved, with an FDR-corrected *p*-value of 0.0356 in the age-limited set.

#### 6. Validation of new TWAS results against existing analyses in ROS/MAP and Mayo cohorts

The number of LOAD-associated genes identified in our analysis of the ROS/MAP cohort (1 763) mirrors recent findings of Canchi et al. (2019)^22^, who performed analyses on 414 ROS/MAP participants and found 1 722 significant genes, albeit with somewhat different pre-processing methods (expression quantified as FPKM rather than expected count), the use of ordinary least squares regression, and the inclusion of *APOE* ε4 status as a regression covariate. To ensure consistency of our ROS/MAP re-analysis with existing work in this cohort, we correlated our association statistics (-log_10_(*p*)) with available published gene-wise effects from two studies (Mostafavi et al. (2018)^23^ and Canchi et al. (2019)^22^) and found strong correlations (r_Canchi_=0.81, r_Mostafavi_=0.86) (**Supplementary Figure 2**).

The large proportion of significant genes (55%) identified in our Mayo cohort TWAS is similar to those identified in published analyses of this dataset^10^ (22 318/55 756 [40%] genes significant in LOAD-control analyses (n=156), not controlling for cell proportions), as well as in the AMP-AD RNAseq reprocessing initiative (Synapse ID: syn14237651; doi: 10.7303/syn9702085) differential expression meta-analysis (7 030/17 013 [41%] significant in Mayo TCX sample subset (n=151)), despite differences in post-quantification quality control and linear modelling pipelines between analyses.

#### 7. Gene-wise comparison of TWAS results across samples

The “tped” metric was calculated as the negative product of gene-wise moderated t-statistics (for the effect of LOAD) calculated from robust linear models. Ranking based on this measure yields a continuous spectrum of concordance, regardless of direction of effect. We used the online REVIGO tool, with default parameters, to summarize many terms that were significant after correction.^24^

In addition to comparing t-statistics between samples analyzed under our unified robust pipeline, we also evaluated correlations between CH and summary statistics from the Accelerating Medicines Project – Alzheimer’s Disease (AMP-AD) consortium reprocessing initiative^18^ (**Supplementary Figure 3**). Specifically, we accessed the LOAD vs. control differential expression results across seven brain regions and three cohorts (ROS/MAP, Mayo cohort, and the Mount Sinai Brain Bank-Mount Sinai School of Medicine (MSBB-MSSM)) processed using different methods (detailed information available at <https://www.synapse.org/#!Synapse:syn9702085>; dio: 10.7303/syn9702085). Again, we identified the strongest correlation between CH LOAD statistics and those from temporal cortex (r= 0.64, C.I._99%_=[0.63, 0.65]), followed by other temporal structures: superior temporal gyrus (r= 0.57, C.I._99%_=[0.56, 0.59]) and parahippocampal gyrus (r= 0.53, C.I._99%_=[0.52, 0.54]) (**Supplementary Figure 3**).

#### 8. Consensus Weighted Gene Co-Expression Network Analysis (WGCNA)

For network construction and co-expression module identification, quality control was again performed on raw count data with two additional steps than for differential expression:1) before removing subjects using PCA and adjusting extreme observations in a gene-wise fashion, counts were log2 transformed (with an offset of 0.5) and 2) outlier genes were identified and removed if their median expression count or count variances were above 1.5 interquartile ranges above the median expression or median variance for all genes, respectively. Before calculating adjacency matrices and deriving co-expression networks, the removeBatchEffect function was used to remove variance due to the same covariates as specified in our differential expression analyses. A design matrix corresponding to AD diagnosis was provided to the removeBatchEffect function to preserve the linear effects of LOAD (i.e. generate partial residuals). Gene adjacency was calculated using signed robust biweight midcorrelations (with the maximum percentile of data that can be considered outliers on either side of the median separately set to 0.05), and a soft thresholding power was selected that maximized network scale free topology based on a topology fit index of at least 0.85 across all three samples (power=12). A deepSplit parameter of 2 was selected for module granularity as this was the point at which the greatest proportion of resulting modules were significantly enriched for biological functions in hypergeometric testing. These criteria have been applied in previous work to benchmark the effectiveness of gene clustering parameters in the context of gene co-expression networks.^25^

#### 9. Cell type proportion estimation from expression data

Human marker genes (n=50 per set) from Darmanis et al. (2015)^26^ were used to estimate proportions of cell types in both samples using a validated singular value decomposition method.^27^

#### 10. Functional enrichment of modules for gene ontology (GO) biological processes using hypergeometric testing

For all three samples, genes assigned to each consensus module were tested for GO term enrichment including categories of biological processes, cellular components, and molecular functions.^28^ Annotations were limited to those including between 10-200 tested genes. GO annotations were extracted from the org.Hs.eg.db and GO.db (version 3.10.0) R packages.^29,30^ Hypergeometric testing was used to evaluate enrichment for GO groups per module, using the full background of genes passing QC for each analysis. To minimize redundancy between these broad categories, we limited our REVIGO^24^-based condensation of GO terms to biological processes only. FDR correction was used (q<0.05) to determine significant enrichment.

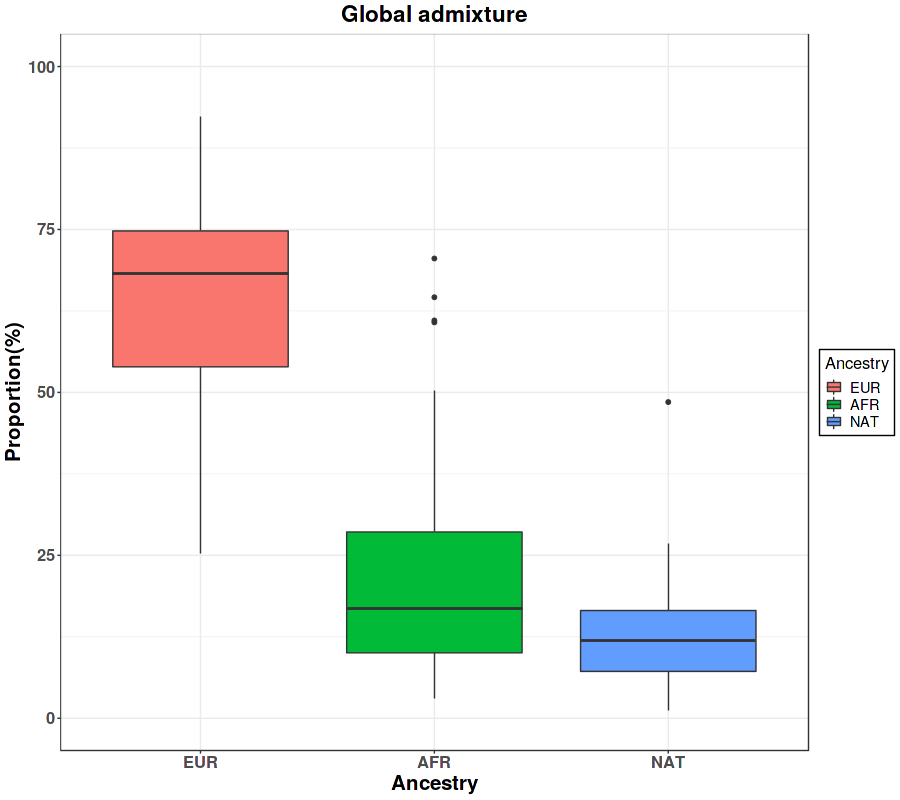

**Supplementary Figure 1**. Global admixture estimation for the CH cohort (EUR=European ancestry; AFR= African ancestry; NAT=Native American ancestry).

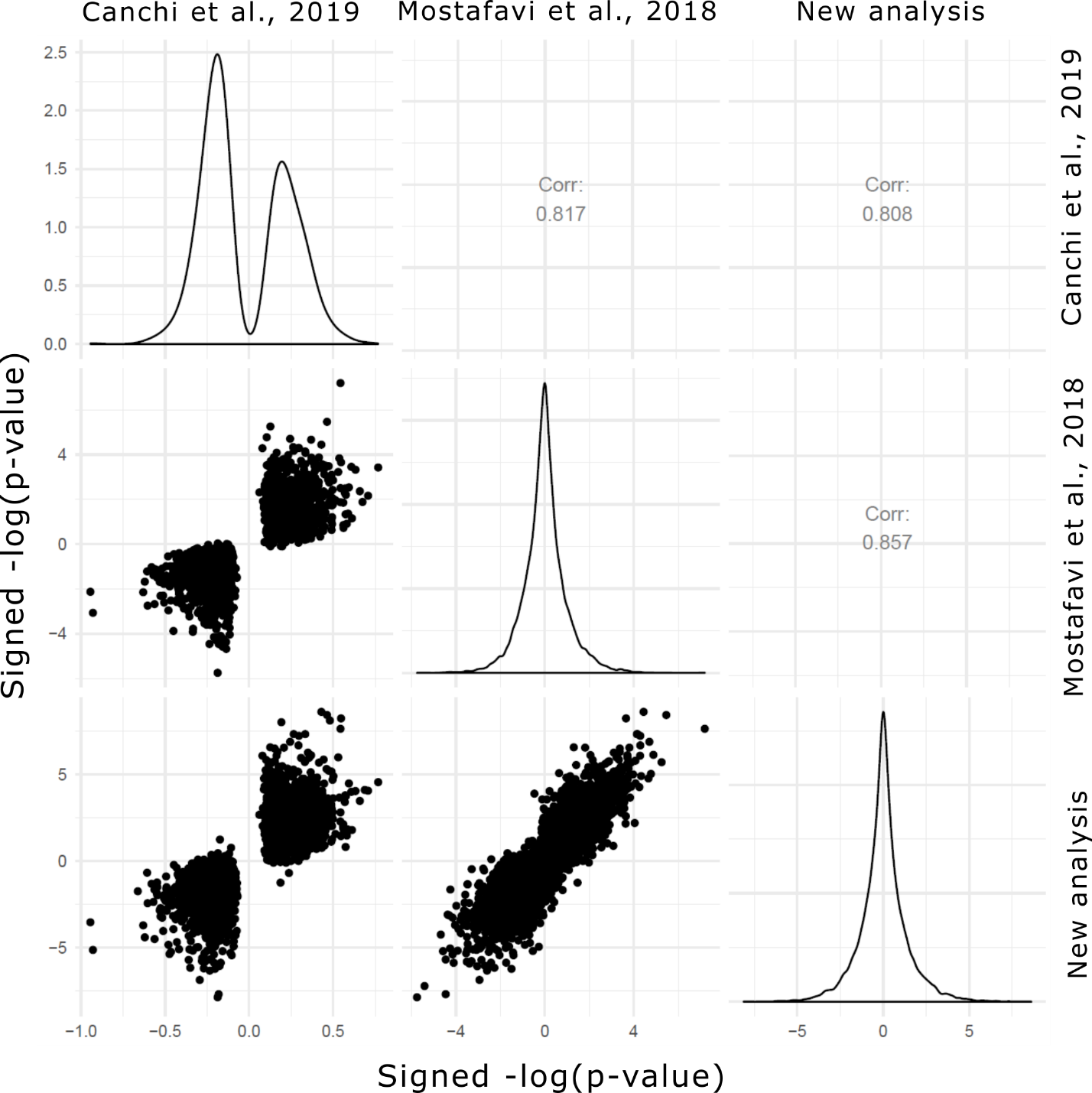

**Supplementary Figure 2**. Comparison of published LOAD vs. non-LOAD differential expression results in the ROS/MAP cohort^22,23^ to our new set of ROS/MAP differential expression results. Due to the availability of summary statistics, signed -log10(p-value) was used as the measure of effect across all three compared samples. Pearson correlation coefficients are shown in the top right quadrant panels, corresponding to its matching panel in the symmetrical matrix.

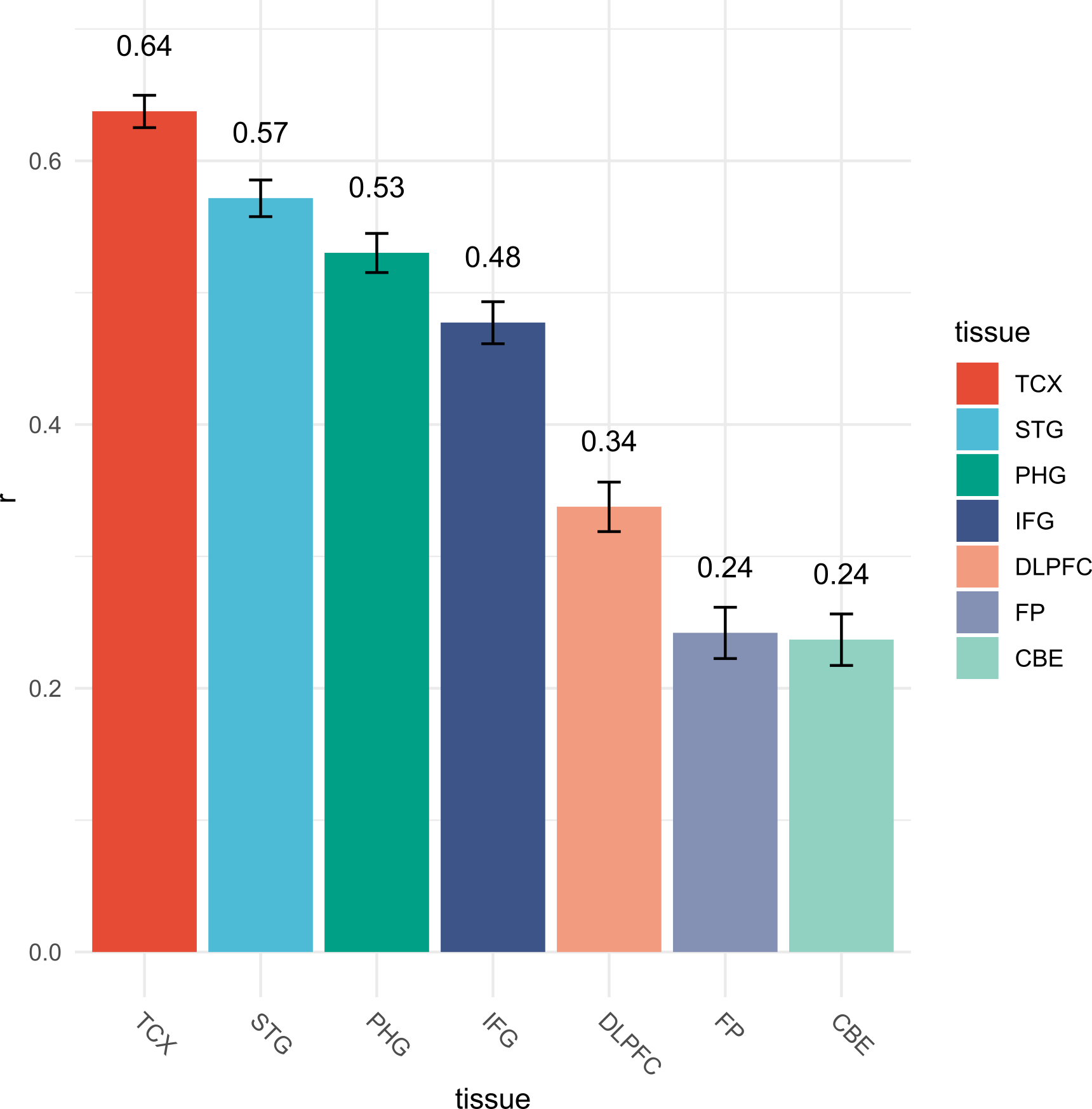

**Supplementary Figure 3**. Pearson correlation coefficients (y-axis, with 99% confidence interval error bars) for the genome-wide correlation of gene-wise LOAD effects (moderated t-statistics) between the CH analysis and those performed in the AMP-AD reprocessing initiative.^18^ Colors represent source tissue region for each analysis. Full summary statistics are available on the Synapse AMP-AD knowledge portal (syn ID: syn17015330). Three separate studies account for the seven tissue regions analyzed: ROS/MAP includes only DLPFC; Mayo Cohort includes TCX and CBE; and Mount Sinai Brain Bank-Mount Sinai School of Medicine (MSBB-MSSM) include STG, PHG, IFG, and FP. For Mayo Cohort and MSBB-MSSM analyses, mixed models were used where the same individual was sequenced in two or more regions. CBE = cerebellum; DLPFC = dorsolateral prefronal cortex; FP = frontal pole (BA 10); IFG = inferior frontal gyrus (BA 44); PHG = parahippocampal gyrus (BA 36); STG = superior temporal gyrus (BA 22); TCX = temporal cortex.

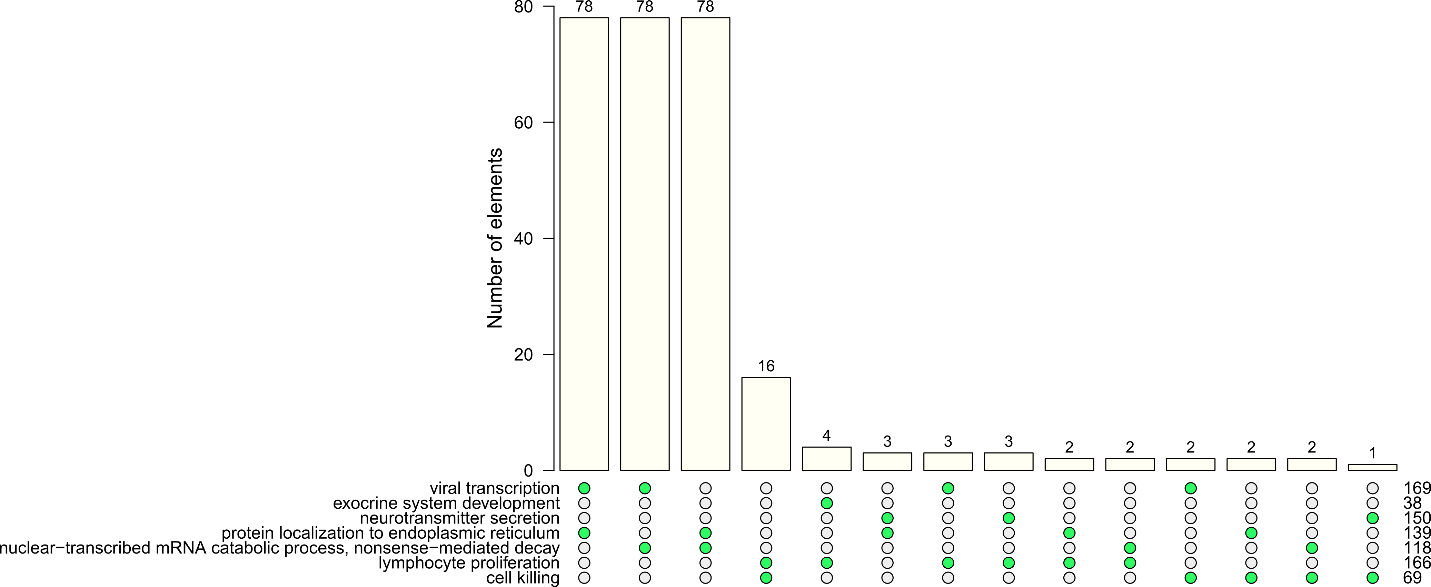

**Supplementary Figure 4**. Overlap of genes belonging to GO groups concordantly and discordantly enriched for differential expression in pairwise comparisons between study samples. “Number of elements” refers to genes. Numbers to the far right of the chart indicate GO group size.
